# Phylogenomic analyses of *Snodgrassella* isolates from honeybees and bumblebees reveals taxonomic and functional diversity

**DOI:** 10.1101/2021.12.10.472130

**Authors:** Luc Cornet, Ilse Cleenwerck, Jessy Praet, Raphaël R. Leonard, Nicolas J. Vereecken, Denis Michez, Guy Smagghe, Denis Baurain, Peter Vandamme

**Affiliations:** BCCM/IHEM, Mycology and Aerobiology, Sciensano, Bruxelles, Belgium; Laboratory of Microbiology and BCCM/LMG Bacteria Collection, Faculty of Sciences, Ghent University, Ghent, Belgium; InBioS–PhytoSYSTEMS, Eukaryotic Phylogenomics, University of Liège, Liège, Belgium; Agroecology Lab, Université libre de Bruxelles (ULB), Boulevard du Triomphe CP 264/02, 1050 Brussels, Belgium; Laboratory of Zoology, Research Institute for Biosciences, University of Mons, Place du Parc 20, 7000 Mons, Belgium; Laboratory of Agrozoology, Faculty of Bioscience Engineering, Ghent University, Ghent, Belgium

**Keywords:** Honeybee, Bumblebee, Microbiome, Metagenomics, *Snodgrassella*, Phylogenomics, Species delineation, Functional analysis, Metabolic modelling

## Abstract

*Snodgrassella* is a Betaproteobacteria genus found in the gut of honeybees (*Apis* spp.) and bumblebees (*Bombus* spp). It is part of a conserved microbiome that is composed of few core phylotypes and is essential for bee health and metabolism. Phylogenomic analyses using whole genome sequences of 75 *Snodgrassella* strains from 4 species of honey bees and 14 species of bumblebees showed that these strains formed a monophyletic lineage within the *Neisseriaceae* family, that *Snodgrassella* isolates from Asian honeybees diverged early on from the other species in their evolution, that isolates from honeybees and bumblebees were well separated and that this genus consists of at least seven species. We propose to formally name two new *Snodgrassella* species that were isolated from bumblebees, i.e. *Snodgrassella gandavensis* sp. nov. and *Snodgrassella communis* sp. nov. Possible evolutionary scenarios for 107 species or group specific genes revealed very limited evidence for horizontal gene transfer. Functional analyses revealed the importance of small proteins, defense mechanisms, amino acid transport and metabolism, inorganic ion transport and metabolism and carbohydrate transport and metabolism among these 107 specific genes.

**Importance:** The microbiome of honeybees (*Apis* spp.) and bumblebees (*Bombus spp*.) is highly conserved and represented by few phylotypes. This simplicity in taxon composition makes the bee’s microbiome an emergent model organism for the study of gut microbial communities. Since the description of the *Snodgrassella* genus, which was isolated from the gut of honeybees and bumblebees in 2013, a single species, i.e. *Snodgrassella alvi*, has been named. Here we demonstrate that this genus is actually composed of at least seven species, two of them (*Snodgrassella gandavensis* sp. nov. and *Snodgrassella communis* sp. nov.) being formally described in the present publication. We also report the presence of 107 genes specific to *Snodgrassella* species, showing notably the importance of small proteins and defense mechanisms in this genus.

**Data summary:**

1. Cornet L and Vandamme P, European Nucleotide Archive (ENA), Project accession: PRJEB47378
2. Cornet L and Vandamme P, European Nucleotide Archive (ENA), Reads accessions: SAMEA9570070 - SAMEA9570078
3. Cornet L and Vandamme P, European Nucleotide Archive (ENA), Genome accessions: GCA_914768015, GCA_914768025, GCA_914768035, GCA_914768045, GCA_914768055, GCA_914768065, GCA_914768075, GCA_914768085, GCA_914768095.

## Introduction

Honeybees (*Apis* spp.) and bumblebees (*Bombus* spp.) harbor a gut microbiome that is important in health and metabolism (1-3). This microbiome is highly conserved with 95% of the gut microbionts falling within a few phylotypes that include Actinobacteria (*Bifidobacterium, Bombiscardovia*), Bacteroidetes (*Apibacter*), Firmicutes (*Lactobacillus* [the so-called Firm-5 or Lacto-1 taxon], *Bombilactobacillus* [Firm-4, Lacto-2) and *Apilactobacillus* [Lacto-3]), Alphaproteobacteria (*Bartonella, Bombella, Commensalibacter*), Betaproteobacteria (*Snodgrassella*) and Gammaproteobacteria (*Frischella, Gilliamella*) (2, 4-18). The genera *Bifidobacterium, Lactobacillus, Bombilactobacillus, Gilliamella* and *Snodgrassella* are generally considered the core microbionts of honeybees and bumblebees (10, 19, 20). These gut-related organisms co-evolved within their hosts during the last 80 million years (9, 10, 21) and contribute to carbohydrate digestion (2, 18, 22) and pathogen defense (23-26). The importance of bees for ecosystem integrity, the contribution of the bee gut microbiota to its hosts’ health, the relative simplicity of the taxonomic composition of the bee gut microbiota, along with their mode of transmission, which is mainly vertical, have made bees an emerging model organism for the study of gut-related microbial communities (1, 18, 27-29).

*Snodgrassella* and *Gilliamella* are physically closely associated within the hind gut, the former growing in contact with the ileum epithelium while the latter forms a dense biofilm on top of the *Snodgrassella* layer, and is in contact with the gut lumen (9, 11, 30). As a result of intricate co-evolution of these two bacteria, horizontal gene transfers (HGTs) have been reported between these two taxa (2, 31). Among others they share Rhs toxin proteins, which are involved in type VI secretion systems (T6SSs) mediated competition (2, 32-34).

The formal description and naming of *Snodgrassella alvi* was based on three strains, one from a honeybee gut sample (wkB2^T^ from *Apis mellifera)* and two from bumblebee gut samples (wkB12 from *Bombus bimaculatus* and wkB29 from *Bombus vagans*) (21). All three strains shared more than 99.1% of their SSU (16S) rRNA gene sequences, which allocated them within the *Neisseriaceae* family. Many additional strains have been reported since, and some phylogenetic analyses revealed a clear separation between *Snodgrassella* isolates of *Apis* spp. and those of *Bombus* spp. (1, 32-35), whereas others reported that the two groups were not monophyletic (10, 22). In the present study we used core gene phylogenomics to demonstrate the separation of *Snodgrassella* isolates into three clades, one colonizing *Bombus* spp. and two colonizing *Apis* spp. We report an early divergence of *Snodgrassella* strains from Asian honeybees, which implies that *Apis*-colonizing strains are paraphyletic. We used average nucleotide identity (ANI) analyses to demonstrate that the genus *Snodgrassella* consists of at least seven species. We further analyzed differential gene contents within these species and examined evolutionary scenarios for the emergence of 107 specific genes. We finally used our own *Snodgrassella* isolates to describe and formally name two of these novel species from bumblebees as *Snodgrassella gandavensis* with LMG 30236^T^ (= CECT 30450^T^) as the type strain and *Snodgrassella communis* with LMG 28360^T^ (= CECT 30451^T^) as the type strain.

## Materials & methods

All custom scripts specifically developed for this study are available at: https://github.com/Lcornet/SNOD.

### Snodgrassella whole genome sequences

Whole genome sequences of 75 *Snodgrassella* isolates were analyzed. To this end, all *Snodgrassella* genomes from the Reference Sequence database of NCBI (RefSeq) (36-37) were downloaded on June 25th, 2021. CheckM v1.1.3 (38) with the *lineage_wf* option was used to identify genomes with contamination levels below 5% and completeness above 95%, which yielded 66 RefSeq genomes. In addition, we determined whole genome sequences of nine bumblebee isolates from an earlier study (15). Together, the genomes of 48 isolates originated from honeybee gut samples (from 4 species) and 27 from bumblebee gut samples (from 14 species). Assembly quality metrics for all 75 genomes were obtained with QUAST v5.0.2 (39). The isolates, along with their geographic origin, and accession numbers, CheckM and QUAST parameters are listed in **Table 1**.

### DNA extraction

Genomic DNA was isolated using a Maxwell® 16 Tissue DNA Purification Kit (cat.# AS1030) and a Maxwell® 16 Instrument (cat.# AS2000). The integrity and purity of the DNA was evaluated on a 1.0% (w/v) agarose gel and by spectrophotometric measurements at 234, 260 and 280 nm. A Quantus fluorometer and a QuantiFluor™ ONE dsDNA System (Promega Corporation, Madison, WI, USA) were used to estimate the DNA concentration.

### Library construction and Genome Sequencing

Library preparation and whole genome sequencing were performed by the Oxford Genomics Centre (University of Oxford, United Kingdom). Library preparation was performed using an adapted protocol of the NEB prep kit. Paired-end sequence reads (PE150) were generated using an Illumina NovaSeq 6000 platform (Illumina Inc., USA).

### Genome assembly

Quality check, trimming of the raw sequence reads and de novo genome assembly were performed using the Shovill v1.1.0 pipeline (https://github.com/tseemann/shovill), which uses SPAdes v3.14.0 (40) as its core and subsamples reads to a sequencing depth of 150x. Contigs shorter than 500 bp were excluded from the final assembly. Quality check of the assembly was performed using QUAST v5.0.2 (39) and CheckM v1.1.3 (38).

### Core gene analysis

ToRQuEMaDA (41), version of June 10th, 2021, was used to select genomes representing the diversity of other *Neisseriaceae* (i.e, bacteria not belonging to the genus *Snodgrassella*). We selected 35 *Neisseriaceae* genomes using the following options of tqmd_cluster.pl: *dist-threshold of 0*.*90, kmer-size of 12 and max-round of 10 and CheckM v1*.*1*.*3 activated* (**Table S1)**. The 110 genomes were then imported into the pangenomic workflow of anvi’o v7.1 (42). The genomes were first loaded individually into anvi’o using the anvi-gen-contigs-database script with default options, and NCBI COGs (43) were associated with each genome-database using the anvi-run-ncbi-cogs script with default options. The consolidated database was constructed using the anvi-gen-genomes-storage script with default options and anvi-pan-genome script with the following options: *min-occurrence of 2 and mcl-inflation of 10*. The protein sequences of all genomes, along with their COG annotations, were extracted from the anvi’o database using the anvi-get-sequences-for-gene-clusters script with default options and the anvi-summarize script with default options, respectively. Functional and geometric indices of anvi’o were computed using the anvi-compute-gene-cluster-homogeneity script with default options. Orthologous groups (OGs) were reconstructed from anvi’o output files using the custom script anvio_pan-to-OGs.py with default options. Finally, core genes were selected using the custom script anvio_OGs-filtration.py with the following options: *pfilter set to yes, fraction set to 1, unwanted orgs limit set to 0, cfilter set to yes, maxcopy set to 1, hfilter set to 1, maxfunctional index set to 0*.*8 and maxgeometricindex set to 0*.*8*. These settings resulted in the selection of 254 core genes out of 11,185 OGs, present in 100 % of the 110 *Neisseriaceae* genomes.

### Phylogenomic analyses

#### *Neisseriaceae* phylogeny

The 254 protein OGs were aligned by anvi’o workflow. Conserved sites were selected using BMGE v1.12 (44) with moderately severe settings (*entropy cut-off* = 0.5, *gap cut-off* = 0.2). A supermatrix of 110 organisms x 85,654 unambiguously aligned amino-acid positions (0.15% missing character states) was generated using SCaFoS v1.30k (45), with default settings. A *Neisseriaceae* phylogenomic analysis was inferred using RAxML v8.1.17 (46) with 100 bootstrap replicates under the PROTGAMMALGF model.

#### *Snodgrassella* phylogeny

*Populibacter corticis* (GCF_001590725.1) was selected as an outgroup for the *Snodgrassella* phylogenomic analysis. The concatenated sequences of 75 *Snodgrassella* and *P. corticis* genomes were extracted from the 254 core gene supermatrix of *Neisseriaceae* to produce a supermatrix of 76 organisms x 85,654 unambiguously aligned amino-acid positions (0.04% missing character states), from which a phylogenomic tree was inferred with RAxML as above. A leave-one-out analysis was then performed by randomly deleting ≈20% of genes present in our dataset. To this end, the corresponding alignments were also reduced to 76 organisms and used to construct one hundred datasets of about 70,000 conserved positions by randomly combining alignment files using the script jack-ali-dir.pl from Bio-MUST-Core (available at https://metacpan.org/dist/Bio-MUST-Core). The 100 supermatrices were assembled using SCaFoS v1.30k (45) with default settings. Trees were inferred using RAxML v8.1.17 (46) using the *fast experimental tree search* method and the PROTGAMMALGF model. A consensus tree was built from the set of 100 trees using the program consense v3.695 (from the PHYLIP package (47), but modified to handle long sequence names), with default settings. In order to test the possible influence of a long branch attraction artefact, the outgroup was eliminated from the supermatrix and a new phylogenomic analysis was performed with the same protocol as above but on a matrix of 75 organisms x 85,654 unambiguously aligned amino-acid positions (0.03% missing character states). We checked the effect of taxon sampling reduction by deleting 40 *Snodgrassella* strains from our supermatrix, while conserving their diversity and the outgroup, using dRep v2.2.3 (48) with default settings. A phylogenetic analysis was inferred with the same protocol as above but using a matrix of 36 organisms x 85,654 unambiguously aligned amino-acid positions (0.22% missing character states).

#### Gut bacteria phylogeny

A total of 486 genomes representing the main phylotypes of bacteria found in the gut of *Apis* spp. and *Bombus* spp. (excluding *Snodgrassella*) were downloaded from RefSeq, prior to filtering the genomes with CheckM v1.1.3 (38) as described above. Also, 247 additional genomes covering a broad diversity of bacteria and archaea were selected with ToRQuEMaDA

(41) using the following options: *dist-threshold of 0*.*90, kmer-size of 12 and max-round of 10 and CheckM activated* (**Table S2**). To perform a large phylogenomic analysis, we used core genes extracted by CheckM. CheckM *taxon set* option was first run to produce a lineage set file. Then the CheckM *analyze* option was run, using the bacterial domain set while providing 843 genomes as input (i.e. 75 *Snodgrassella* genomes, 35 *Neisseriaceae* genomes and the 733 genomes mentioned above). The CheckM *qa* option was then used to produce the marker files. The custom script Checkm-to-OGs.py allowed to reconstruct OGs from CheckM marker files, selecting only unicopy OGs, thereby producing 76 OGs. The OGs were aligned using MAFFT v7.453 (49), run with the *anysymbol, auto and reorder* parameters. Conserved sites were selected using BMGE v1.12 (44) with moderately severe settings (*entropy cut-off* = 0.5, *gap cut-off* = 0.2). A matrix of 843 organisms x 12,930 unambiguously aligned amino-acid positions (6.72 % missing character states) was produced using SCaFoS v1.30k (45), with default settings. A phylogenomic tree was produced with the same protocol as above for large phylogenomics.

#### Average nucleotide identity analysis

Pairwise average nucleotide identities were determined for 75 *Snodgrassella* genomes and the *P. corticis* (GCF_001590725.1) genome using fastANI v1.32 (50) with default settings.

#### Specific gene analysis

A new anvi’o pangenomic dataset using 75 genomes of *Snodgrassella* and *P. corticis* (GCF_001590725.1) was constructed using the same protocol as described above. The custom script anvio_OGs-filtration.py was used to determine which genes occurred specifically in a given group of organisms. We used the same options as for the core genes, with the exception that the number of representatives of a group was set to 0.6 (60%). The inflation number was set to 10 in anvi’o (42), and the value of 60% recovered orthologous groups even if the inflation parameter was too strict for some of them. All members of a group had to comprise an OG before it was considered specific. An orthologous enrichment was then performed to compensate for the potential lack of sensitivity in orthology inference. In addition, to enrich the OGs with potentially absent members of the group due to the 60% threshold, the enrichment could also add any *Snodgrassella* from other groups to control that the added sequences were indeed specific and not the result of a false orthology inference. We performed the orthologous enrichment with Forty-Two v0210570 (51-52) (available at https://metacpan.org/dist/Bio-MUST-Apps-FortyTwo). We used the 843 genomes that included the 75 *Snodgrassella* genomes to perform the enrichment. The custom script Confirm-OGs.py was used to validate the specific OGs based on the criteria that all *Snodgrassella* of the considered group, but no *Snodgrassella* from other groups, must be present in the OGs after enrichment. We performed this approach on all groups defined in the present study, as well as all pairwise combinations of them, including one with all groups together (i.e, all S*nodgrassella* strains). All nodes of the tree have also been tested to assess the presence of specific genes shared between groups defined in this study. A total of 107 individual gene phylogenies were computed by aligning the sequences using MAFFT v7.453 (49), with the same options as above, selecting the conserved sites with BMGE v1.12 (44), with the same options as above, and inferring the trees with RAxML v8.1.17 (46), with 100 bootstrap replicates under the PROTGAMMALGF model.

#### Metabolic analyses

The functions of the group-specific OGs were first determined using NCBI COGs (43) if the information was available as a product of the anvi’o workflow. COG pathways and COG functions were used to label the metabolic functions. When COG analyses yielded no results, Mantis (53) was used with default settings, and the information from the consensus annotation file was used to label function. OGs without any hit in COG or Mantis analyses were considered as unknown genes while OGs having hits with proteins of unknown function were classified as “Function unknown”.

#### Horizontal gene transfer

GTDBtk (54) was used to infer the GTDB taxonomy (55) of the 561 phylotype genomes (75 *Snodgrassella* and 486 representatives of the other bacterial bee gut phylotypes; see above). MetaCHIP v1.10.15 (56) with default settings was then used to infer HGT. The group-specific genes were then searched in the Metachip results using BLASTP v2.2.28 (57) with a threshold of 98% of identity on 95% of query length.

#### Biochemical characterization

Biochemical characteristics were determined for the bumblebee isolates LMG 28360^T^, R-53528, R-54236, LMG 30236^T^ and R-53680, and for the type strain of *S. alvi* NCIMB 14803^T^. Growth was tested on nutrient agar (Oxoid), TSA (Oxoid), BHI-agar (BD Difco), Columbia Agar (Oxoid) supplemented with 5% sheep blood and All Culture (AC) agar (2% tryptose, 0.3% beef extract, 0.3% yeast extract, 0.3% malt extract, 0.5% dextrose, 0.02% ascorbic acid, 1.5% agar; w/v)) after two, three and four days of incubation at 37°C in a CO_2_ enriched atmosphere (6% CO_2_, 15% O_2_) provided by CO_2_Gen Compact sachets (Oxoid). Hemolysis of sheep blood was checked. Growth was also tested at 37 °C in ambient atmosphere (0.04% CO_2_, 20.95% O_2_) on AC agar, and in anaerobic atmosphere (9-13% CO_2_, <1% O_2_) provided by AnaeroGen sachets (Oxoid) on AC agar and on AC agar supplemented with 10 mM KNO_3_. For the tests mentioned below, cultures were incubated in a CO_2_ enriched atmosphere. The temperature growth range was tested after two days of incubation on AC agar at 4, 15, 20, 28, 37, 40 and 45 °C. Cell and colony morphology, motility, oxidase and catalase activities and Gram stain reaction were assessed on cultures grown for two days on AC agar at 37°C. Motility was determined by examining wet mounts in broth by phase contrast microscopy. Oxidase and catalase activities and Gram staining were tested using conventional procedures (58). The effect of NaCl on growth was investigated in AC broth supplemented with different concentrations of NaCl (0–10% with 1% intervals, w/v) after 3 days of incubation. The pH range for growth was evaluated after 3 days of incubation in AC broth buffered at pH 4.0 to 9.0 at intervals of 1 pH unit using the following buffer systems: acetate buffer (4.0–5.0), phosphate buffer (pH 6.0– 8.0) and Tris–HCl (pH 9.0). Nitrate reduction and denitrification, and urease, indol and H2S production were tested using standard microbiological procedures (58) after 72h of incubation.

DNase agar (Sigma-Aldrich), AC agar with 0.8% gelatin (Merck, w/v), AC agar with 0.8% soluble starch (w/v), AC agar with 1.3% dried skim milk (Oxoid, w/v) and AC agar with 1% tween 20 or 80 (v/v) and were checked for growth after 72 of incubation and for DNase activity, gelatinase activity, hydrolysis of starch, casein, tween 20 and 80, respectively.

## Results and discussion

### Species delimitation within the *Snodgrassella* genus

We studied the diversity within the genus *Snodgrassella* using extensive phylogenomic and average nucleotide identity (ANI) analyses (**Figure 1**). Highly conserved genes were selected to perform the phylogenomic analyses. Multiple events of HGT affecting *Snodgrassella* genomes have been reported (2, 31-32) and such events may be damaging for inferring species phylogenies (59-61). To minimize interference of HGT, we used shared *Neisseriaceae* core genes only, by incorporating 35 non-*Snodgrassella* bacteria from this family into our dataset. We further imposed a strict unicopy presence to core genes in order to avoid artifacts linked to paralogous sequences, which also have been reported to be deleterious to phylogenomic studies (62-63). Finally, we enforced high geometric and functional indices (0.8 for both), resulting in the selection of genes with less than 20% of gap insertions and 20% of substitutions, respectively. Our phylogenomic dataset was thus composed of 254 genes that were structurally and functionally conserved within the family *Neisseriaceae*. Our analyses of the *Neisseriaceae* phylogeny confirmed with 100% bootstrap support that the genus *Snodgrassella* is monophyletic within the family *Neisseriaceae* (**Supplemental Figure 1**) (21) (64). Its nearest neighbor taxon was *P. corticis* (63). Prior to the description of *P. corticis* in 2017 (64), the lineage formed by *Stenoxybacter acetivorans* and *Snodgrassella* spp. was suggested to represent a gut-specific clade within the Betaproteobacteria, based on the observation that also *S. acetivorans* was isolated from an insect, i.e. termite, gut sample (21). The description of *P. corticis*, an organism isolated from bark tissue of poplar canker, appeared to disrupt this image, yet, more should be known about the latter organism before the hypothesis of a gut-specific clade within the Betaproteobacteria is abandoned. In the rest of the present study, we used *P. corticis* CFCC 13594^T^ as an outgroup for studying *Snodgrassella* phylogeny.

**Figure 1.**
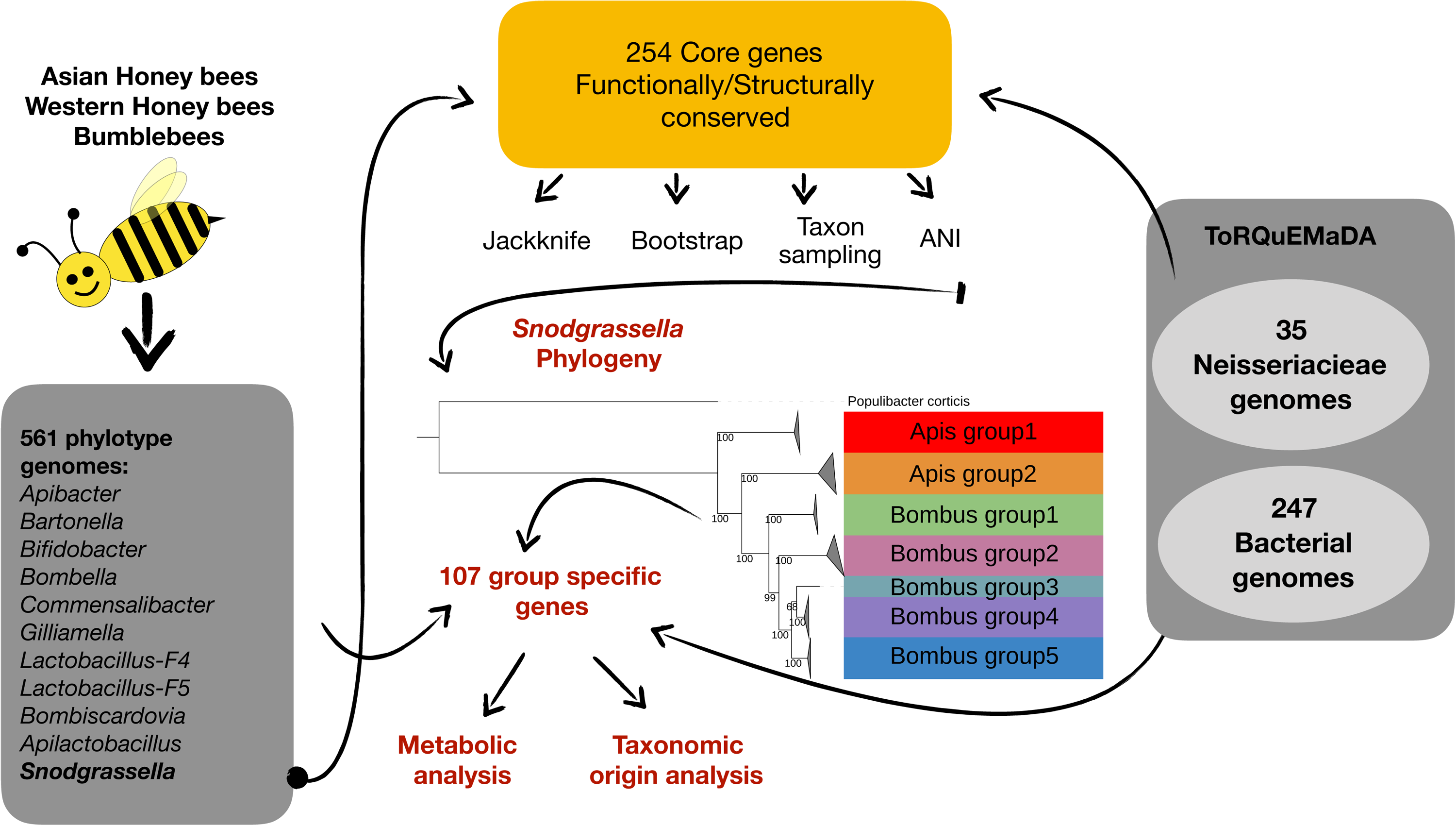
Graphical abstract. We used 9 newly sequenced *Snodgrassella* genomes and 67 (including the outgroup) public genome assemblies to reconstruct the phylogeny of the genus *Snodgrassella* using 254 core genes, all functionally and structurally conserved. We used independent phylogenetic methods (bootstrapping, leave-one-out and taxon sampling) to assess the robustness of our phylogenomic trees. Combined with an Average Nucleotide Identity analyses, our results indicated new species delimitations within this genus. We further investigated specific gene content within the species groups, and their functional importance.

Our phylogenomic and leave-on-out analyses both revealed a clear separation between *Snodgrassella* isolates from *A. mellifera* and *Bombus* spp., with full support values in bootstrap and leave-one-out analyses (**Supplemental Figure 2**). The 75 *Snodgrassella* strains formed three clades. Those isolated from Asian honeybees, referred to as Apis group1, formed a monophyletic group separated from the two other clades with full support in both bootstrap and leave-one-out analyses (**Supplemental Figure 2**). The same three clades were reported by Powell et al. (2016) (65) using *minD* as a single phylogenetic marker gene. Yet, the latter study did not use an outgroup to root the obtained phylogeny. Using *P. corticis* CFCC 13594^T^ as an outgroup, we demonstrated that Apis group1 is the first diverging group of the *Snodgrassella* phylogeny (**Supplemental Figure 2**). The separation into three lineages was confirmed by removing *P. corticis* CFCC 13594^T^ from the phylogenomic analyses in order to check the presence of a long branch attraction artefact (**Supplemental Figure 3**). The *Snodgrassella* phylogeny showed shorter branch lengths in *Snodgrassella* isolates from *Apis mellifera* (hereafter referred to as Apis group2) than in branches with isolates from *Bombus* spp. (**Figure 2**). The latter represented five lineages (**Figure 2, Table 1**). The first, Bombus group1, was composed of four isolates from *Bombus pensylvanicus* (all collected in the USA) and was fully supported by the bootstrap and leave-one-out analyses (**Supplemental Figure 2**). The second, Bombus group2, consisted of a single isolate from *Bombus nevadensis* (USA). The three remaining groups were fully supported, in the bootstrap and leave-one-out analyses, as well, and were referred to as Bombus group3, which was composed of three isolates from *Bombus appositus* (USA), Bombus group4, which was composed of two isolates from *Bombus pascuorum* and *Bombus lapidarius* (Belgium), and Bombus group5, which comprised 17 isolates collected from 11 *Bombus* species in Belgium and the USA (**Figure 2, Table 1**) and which included the isolates wkB12 and wkB29, reported by Kwong and Moran (21). This topology was confirmed with sparser taxon sampling, after removing 40 strains with dRep (48) and computing another large phylogenomic analysis (tree of **Figure 3)**.

**Figure 2.**
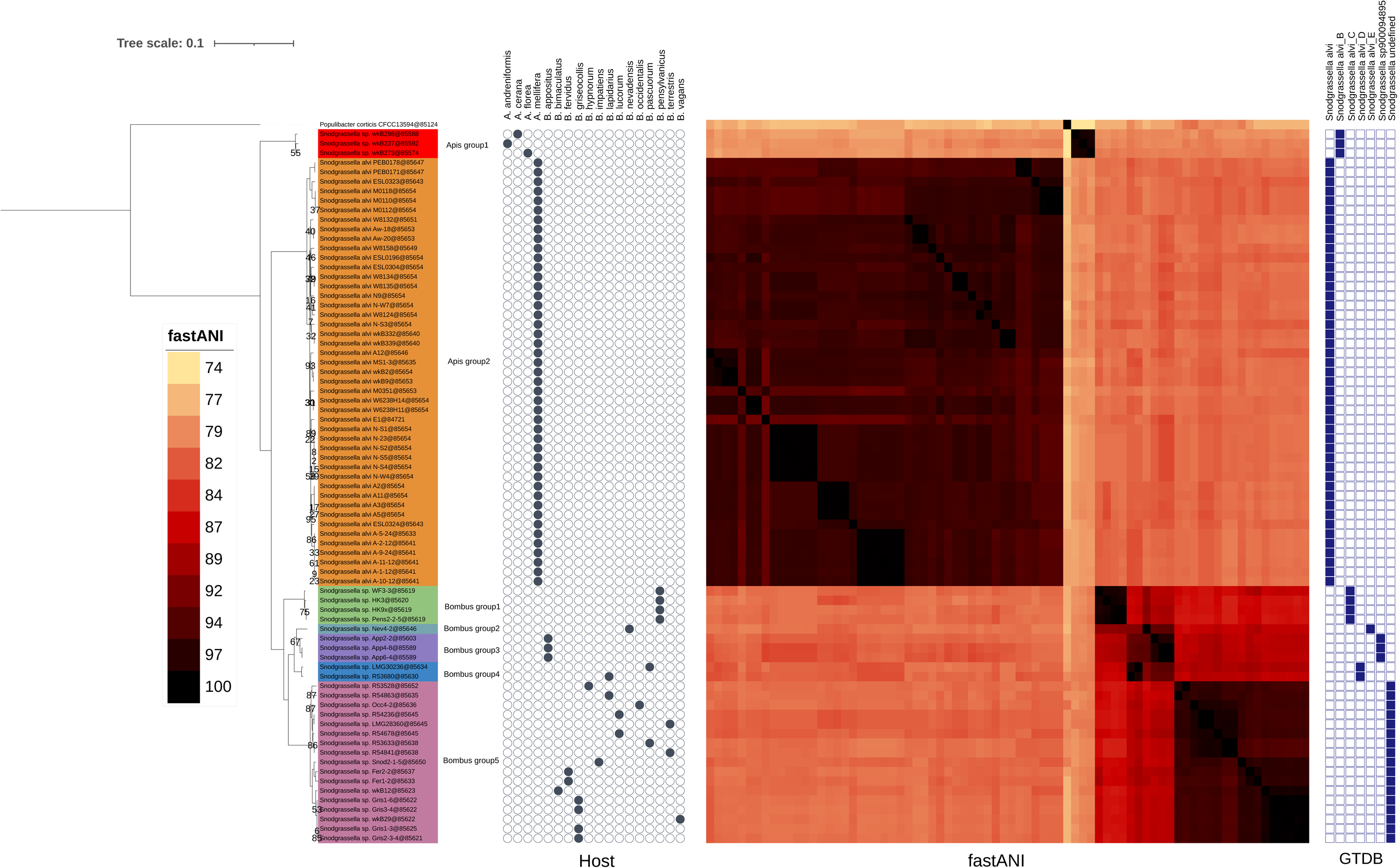
*Snodgrassella* phylogeny and ANI comparison. A maximum likelihood tree was inferred from 254 core gene sequences under the PROTGAMMALGF model with RAxML (44) from a supermatrix of 76 organisms x 86,654 unambiguously aligned amino-acid positions. Only bootstrap values below 100 are shown at the nodes. ANI values, indicated under the heatmap legend, were computed with fastANI (48). Host origins were taken from biosample metadata in the NCBI portal, except for the 9 newly sequenced genomes, for which the metadata were taken from reference (15). GTDB hits, blue squares on the right, were determined using GTDBtk (53).

**Figure 3.**
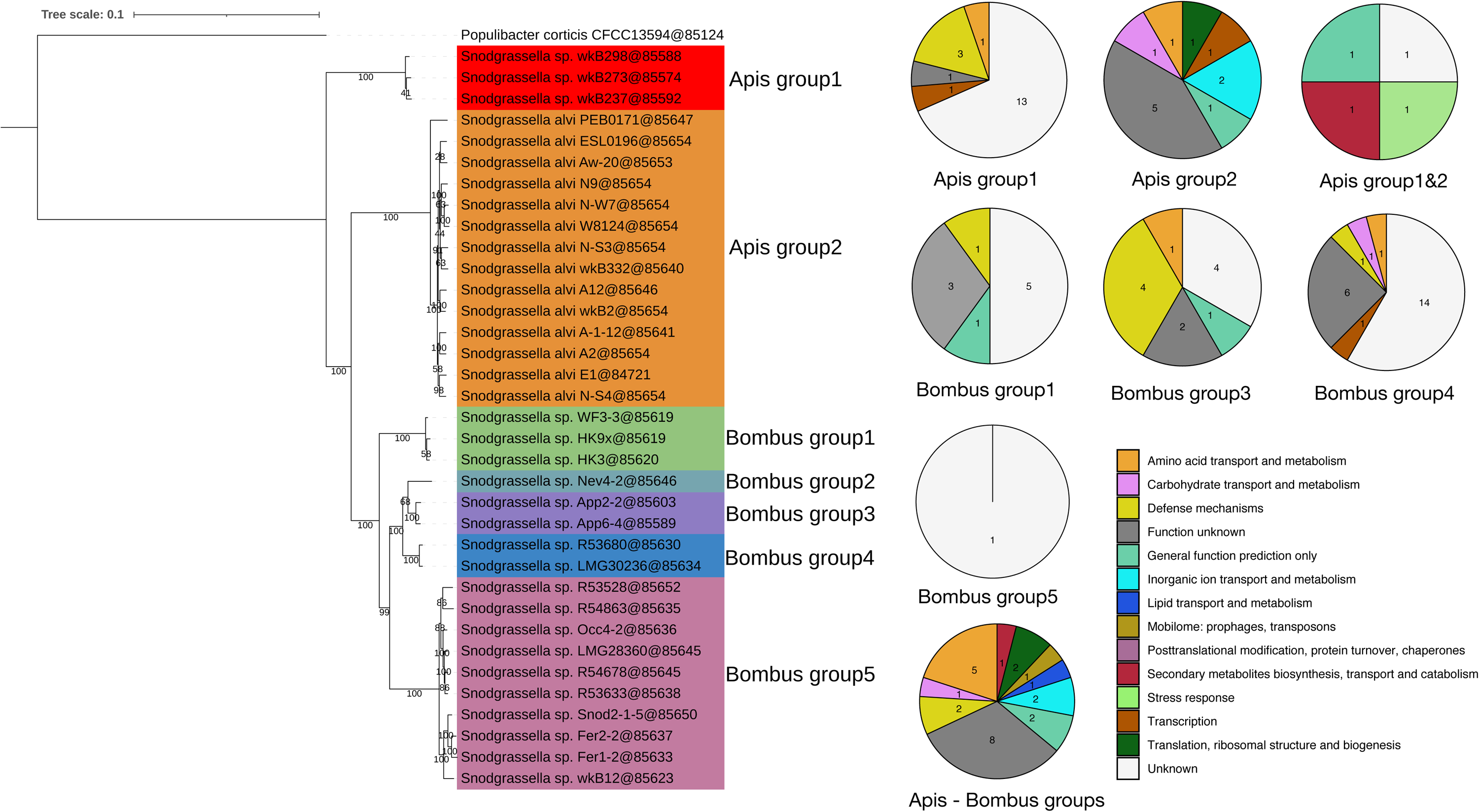
*Snodgrassella* phylogeny, after dereplication of highly similar genomes, and metabolic analysis per species group. The maximum likelihood tree was inferred on 254 core genes under the PROTGAMMALGF model with RAxML (44) from a supermatrix of 36 organisms x 86,654 unambiguously aligned amino-acid positions. Functional analyses were performed using COG (41) and Mantis (52). Numbers indicated in the pie-charts correspond to absolute OGs numbers. Specific genes were computed for entire groups, before dereplication. Bootstrap support values are shown at the nodes.

To quantify the taxonomic divergence between the observed phylogenomic groups we performed ANI analyses. The latter are generally used for species delimitation in bacterial taxonomy (66) with ANI values of about 95-96% corresponding to the species delineation threshold (67-68). The ANI analysis (**Figure 2, part ANI**) revealed values within each of the phylogenomic groups (except Bombus group2, for which only a single genome was available) that were consistently above 95% ANI, while ANI values between genome sequences of different groups were consistently below 87%. Therefore, each of the seven phylogenomic groups corresponded with a distinct *Snodgrassella* species in modern bacterial taxonomy. Together with *S. alvi*, these seven species also corresponded with the seven *Snodgrassella* species defined in the GTDB (https://gtdb.ecogenomic.org/searches?s=al&q=Snodgrassella) database (55) (**Figure 2, GTDB part**). The *S. alvi* type strain wkB2^T^ clustered within Apis group2 and by taxonomic convention, the name *S. alvi* should therefore be restricted to organisms from Apis group2. All other phylogenomic groups detected in the present and in earlier studies represent novel *Snodgrassella* species. Below we propose to formally name Bombus group4 as *Snodgrassella gandavensis* with LMG 30236^T^ (= CECT 30450^T^) as the type strain and Bombus group5 as *Snodgrassella communis* with LMG 28360^T^ (= CECT 30451^T^) as the type strain. The formal naming of the remaining phylogenomic groups, i.e. Apis group1, Bombus group1, Bombus group2 and Bombus group3, can be done upon characterization and public deposit of reference cultures conform to standard practices in bacterial taxonomy (66).

It has been suggested that the transmission mode of *Snodgrassella* between workers and larvae is mainly vertical within a colony (27, 32, 69). Yet, Koch et al. (35) demonstrated exchange of strains between hosts of different colonies, thus rejecting strict co-speciation. Although for most of the groups, i.e. *Snodgrassella* species, detected in the present study only a limited number of isolates or genomes was available, our data showed that indeed several *Snodgrassella* species occur in more than one *Bombus* species and therefore reinforced the rejection of strict co-speciation. The close phylogenetic relationship between *Snodgrassella* species associated with bumblebees and *Snodgrassella* species associated with the Western honeybee additionally supports the hypothesis of horizontal transfer of symbionts between bee clades. The origin of extant bumblebees is estimated to the Miocene, as based on fossil records and molecular phylogeny (70-71). The origin of this clade is associated with a global cooling of the Palearctic region (72). Ancestors of bumblebees were likely sharing host plants with the Western honeybee ancestor clade which is also associated with a temperate climate, and not with the *Apis cerana* honeybee clade which is associated with a subtropical to tropical warmer climate. Overall, the similarity of microbiota among corbiculate bees seems to be related to both phylogeny and sharing of common habitats/climate.

### Specific gene analysis

*Snodgrassella* has been co-evolving with other bacteria in the gut of honeybees and bumblebees for 80 million years (9, 10, 21). Interestingly the age of the common ancestor of the clade of the hosts (i.e. corbiculate bees which include both bumblebees and honeybees) is also estimated round ∼85 millions years (73). Different species and genotypes have evolved and may have developed different functions in their hosts. We investigated the presence of group specific genes (i.e., genes that are exclusively present in one group [or species] and that are shared by all members of this group) and determined their functionalities. The rationale was that a conserved gene probably confers a benefit to members of that group. We detected 107 specific genes: 19 were specific for Apis group1, 12 for Apis group2 (i.e. *S. alvei*), 10 for Bombus group1, 12 for Bombus group3, 24 for Bombus group4 (*S. gandavensis*) and 1 for Bombus group5 (*S. communis*) (**Figure 3**). No specific genes were found for Bombus group2. In a next step, genes shared by multiple groups or species were examined. Only two group combinations presented specific genes: Apis group1 and Apis group2 shared 4 genes that were absent in all others, and all *Snodgrassella* together shared 25 genes (**Figure 3**).

Of these 107 genes, 37 did not correspond to proteins known in COG or Mantis (53). The latter is the most recent and comprehensive tool for protein annotation and contains Pfam, eggnog, NPFM, TIGRfams and Kofam family data for the three domains of life (53). These 37 proteins were relatively short (median = 53 aa; IQR = 46) compared to the rest of the 107 genes (median = 197 aa; IQR = 202.25) with only six proteins above 100 aa (Table 2). Small proteins in bacteria can have roles in transport, signal transduction, act as chaperones, be involved in stress responses or virulence (74-75) and can also be used as bacteriocins (76). Unknown genes are a recurrent issue in metagenomic studies where they may represent up to 40% of the genes (77). Beside these genes without match, 24 genes encoded proteins with unknown function but with hits in COG or Mantis, while six genes had a general predicted function only and could not be affiliated with a metabolic pathway. Together, 67 of 107 specific genes (62.6%) and their putative benefits remained unknown (Table 2, **Figure 3**).

Eleven of the 40 identified specific genes represented defense mechanisms, which are important for gut colonization in bees (14) (Table 2, **Figure 3**). The two most conserved defense mechanisms (4 genes each) corresponded to drug exporter and Rhs proteins (**Table 2**). These two defense mechanisms generate similar benefits and can provide competitive advantages. Steele and Moran (34) recently demonstrated that Rhs proteins represent different toxins, which are injected in neighboring cells through a type VI secretion system T6SS (2, 32-34). The four Rhs proteins found in the present study were specific genes of Bombus group3, an endosymbiont thus far detected only in *B. appositus*. The presence of specific toxins in these three strains might explain the specificity for their host by T6SS mediated-competition. The last three defense mechanisms included proteins for glycosylation of phage DNA (Bombus group1), sensing of invasive viruses and antibiotic production (Apis group1) (**Table 2**). Apis group1 possessed a specific protein, closely related to the 2-oxo-3-(phosphooxy)propyl 3-oxoalkanoate synthase, which triggers the production of the antibiotic virginiamycin in *Streptomyces virginiae* (78).

Among the remaining identified specific genes, 9 genes belonged to the metabolic category amino acid transport and metabolism, 4 genes belonged to inorganic ion transport and metabolism and 4 genes belonged to carbohydrate transport and metabolism (**Figure 3**). Because host gut environments are scarce in iron and amino acids, de novo biosynthetic pathways and iron importers have been reported to be essential for gut host colonization (79). Specific genes linked to carbohydrate metabolism might provide a benefit linked to a carbohydrate-rich diet (1), even if such genes are more commonly found in *Gilliamella* than *Snodgrassella* genomes (30). The 12 remaining identified specific genes represented different metabolic pathways with unclear function in, or benefit to, the host (**Table 2)**.

### Horizontal gene transfer

Horizontal gene transfer is a process that plays a major role in bacterial evolution. Because of the spatial proximity of *Snodgrassella* and other gut bacteria during 80 millions years of coevolution, HGT events may have impacted specific gene complement. We investigated the potential presence of such events by using Metachip (56). We included all bee gut symbiont genomes available in RefSeq (561 genomes, i.e. 75 *Snodgrassella* + 486 representatives of other bee gut symbionts) to maximize the odds of HGT detection and detected only 57 events of HGT in which *Snodgrassella* spp. were donor or recipient (**supplemental Figure 4**, Table S3). This is less than the 87 events reported by Kwong et al. (2). Metachip is designed to infer HGT from genomes assembled from microbial communities (56) and detects potential HGT events using a BLAST best-hit approach, and then validates these hits by performing a duplication-transfer-loss (DTL) reconciliation between a species tree and an individual gene tree. The DTL filter used by Metachip is more stringent than the BLAST e-value threshold of 1-e50 used by Kwong et al. (2), which may explain this difference.

None of the 107 specific genes described above were present among the 57 HGT affected genes detected by Metachip using BLASTP and a filter on 98% identity (**supplemental Table 3**). We investigated the absence of HGT in the group of 107 specific genes further by enriching the comparison with orthologous sequences taken from the 561 gut symbiont genomes used above and with orthologous sequences from the 247 genomes unrelated to the bee gut ecosystem, thereby covering also bacterial and archaeal diversity (**Figure 4**). This allowed us to analyze the taxonomic diversity of the specific genes and revealed that 68 genes were unique to *Snodgrassella* while 39 genes were found in other bacteria too (27 genes in multiple other bacteria and 12 genes in only one other bacteria) (**Figure 4**). The evolutionary history of the 68 genes unique to *Snodgrassella* was explained more easily by duplication in *Snodgrassella* than by acquisition by HGT or gene loss in all other bacteria. In contrast, the 27 specific genes found in other bee endosymbionts or in other bacteria unrelated to the bee gut microbiome suggest that gene loss in other *Neisseriaceae* was the most plausible evolutionary path of the genes (**Figure 4**). We computed individual gene trees for these 27 OGs (available at https://github.com/Lcornet/SNOD) and compared them manually to species trees (**supplemental Figure 5**). For only 8 genes out of the 27 (indicated in **Table 2**), *Snodgrassella* sequences clustered with those of other bee endosymbionts, which may suggest HGT events. For 12 genes (out of 107), no convincing evidence of HGT was detected because these genes were only present in two species, making the comparison to the species tree impossible. Ten of the latter genes were detected in *Gilliamella* and *Snodgrassella* only, and two were detected in *Snodgrassella* and *Frischella* only. These genes included three out of four specific Rhs proteins of Bombus group3, which were shared with *Gilliamella* genomes.

**Figure 4.**
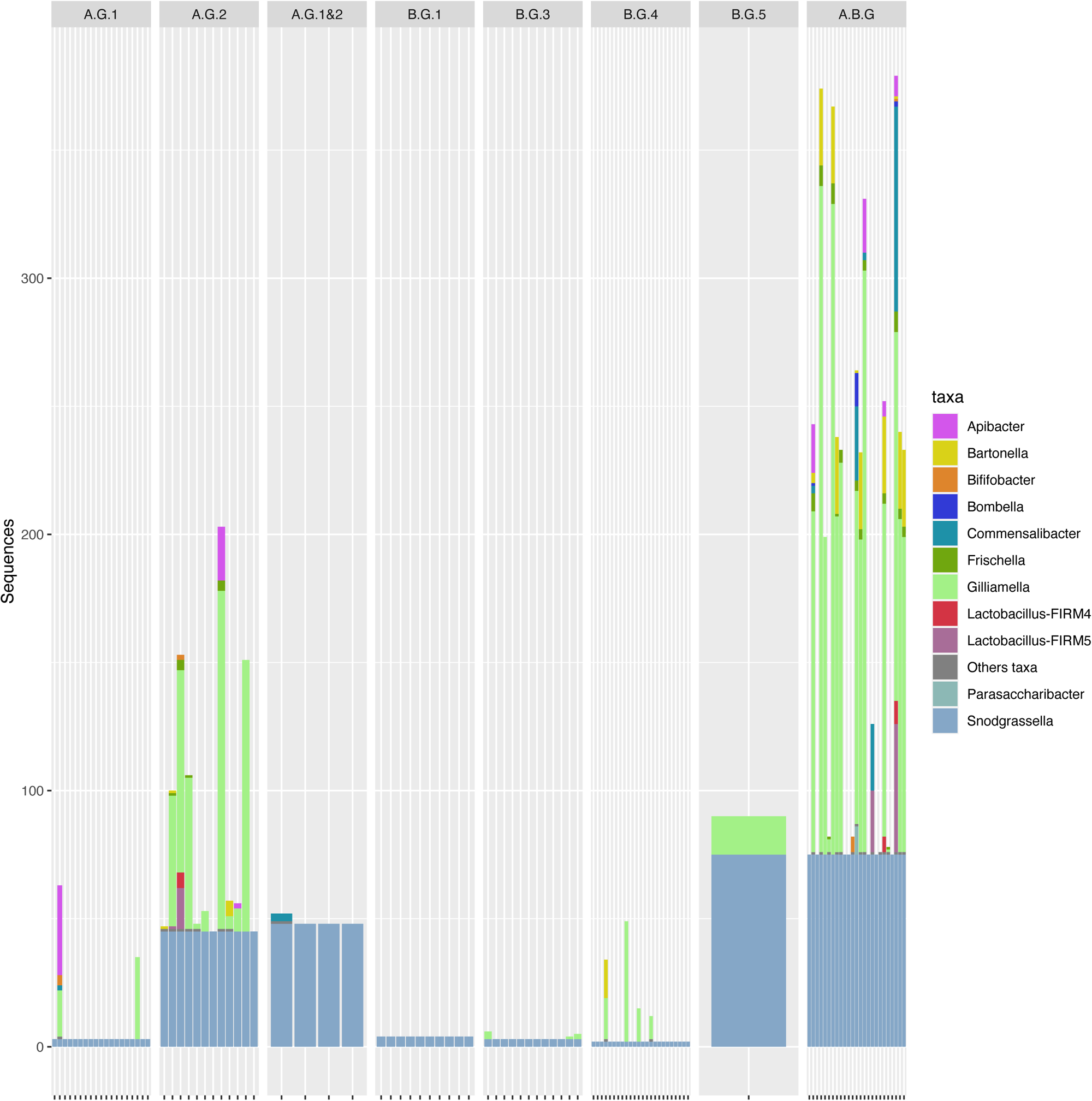
Taxonomic profile of group-specific genes. 107 specific genes from bee gut bacteria (incl. 75 *Snodgrassella* genomes) were enriched with sequences from 486 genomes from bee gut and 247 genomes from bacteria not related to the gut of bees, using Forty-Two (50-51). The Y axis represents the absolute number of sequences and the X axis represents individual genes. Genes were grouped by species group: A.G.1 for Apis group1 (19 genes), A.G.2 for Apis group2 (12 genes), A.G.1&2 for the two Apis groups together (4 genes), B.G.1 for Bombus group1 (10 genes), B.G.3 for Bombus group3 (12 genes), B.G.4 for Bombus group4 (24 genes), B.G.5 for Bombus group5 (1 gene) and A.B.G for Apis and Bombus groups together (25 genes).

## Conclusion

In the present study we used highly conserved *Neisseriaceae* core genes to perform multiple phylogenomic analyses. We demonstrated the monophyly of *Snodgrassella* strains isolated from bumblebees and the paraphyly of strains isolated from honeybees. We also demonstrated that *Snodgrassella* strains from Asian honeybees are an early diverging group. Combined with average nucleotide identity analyses, our data further indicated that this genus comprises at least seven species. Below we describe and formally name two new *Snodgrassella* species from bumblebees, i.e. *S. gandavensis* sp. nov. for Bombus group4 (with LMG 30236^T^ as the type strain) and *S. communis* sp. nov. for Bombus group5 (with LMG 28360^T^ as the type strain). Additional information is provided in the supplementary information. We detected 107 specific genes among these seven species. For the large majority of these 107 genes there was no evidence for HGT. Functional analyses revealed the importance of small proteins, defense mechanisms, amino acid transport and metabolism, inorganic ion transport and metabolism and carbohydrate transport and metabolism among these specific genes.

### Description of *Snodgrassella gandavensis* sp. nov

*Snodgrassella gandavensis* (gan.da.ven’sis. M.L. masc. adj. *gandavensis* of *Gandavum*, the Latin name for Ghent, referring to the place where these bacteria were first isolated).

Cells are non-motile, Gram stain-negative rods, about 1.2 µm long and 0.8 µm wide that occur singly or in pairs. Optimal growth on rich agar media at 37°C in a CO_2_ enriched atmosphere. Grows in anaerobic conditions when supplemented with 10 mM KNO_3_, but not in aerobic conditions. Positive for catalase activity and nitrate reduction. Negative for oxidase activity. The type strain is LMG 30236^T^ (= CECT 30450^T^), which was isolated in 2015 from the gut of *Bombus pascuorum* sampled in 2015 in Wetteren, Belgium. The whole genome sequence of LMG 30236^T^ has a size of 2.51 Mbp. The DNA G+C content is 43.76 mol%. The whole genome sequence is publicly available under accession number GCA_914768025.1. The 16S rRNA gene sequence is publicly available under accession number OU943324.

### Description of *Snodgrassella communis* sp. nov

*Snodgrassella communis* (com.mu’nis. L. fem. adj. *communis* common, because of its wide host range).

Cells are non-motile, Gram stain-negative rods, about 1.2 µm long and 0.8 µm wide that occur singly or in pairs. Optimal growth on rich agar media at 37°C in a CO_2_ enriched atmosphere. Grows in anaerobic conditions when supplemented with 10 mM KNO_3_. Growth in aerobic conditions is strain dependent. Positive for catalase activity and nitrate reduction. Negative for oxidase activity.

The type strain is LMG 28360T (= CECT 30451T), which was isolated in 2013 from the gut of *Bombus terrestris* sampled in 2013 in Ghent, Belgium. The whole genome sequence of LMG 28360^T^ has a size of 2.31 Mbp. The DNA G+C content is 43.26 mol%. The whole genome sequence is publicly available under accession number GCA_914068745.1. The 16S rRNA gene sequence is publicly available under accession number OU943323.

## Authors contributions

LC, IC, DB, PV conceived the study. IC assembled the newly sequenced genomes. RRL carried out the ToRQuEMaDA analyses. LC performed the rest of the analyses and drew the figures. LC, IC, DM, DB, PV wrote the manuscript. PV, DB, NJV, DM and GS provided part of the funding. All authors read and revised the manuscript.

## Acknowledgments

The authors thank David Colignon and the CÉCI for their help with computing cluster usage, and Cindy Snauwaert for performing the phenotypic characterization tests.

## Figures and Tables

**Table 1. Details of the *Snodgrassella* strains and public assemblies**. Completeness and contamination values/metrics were estimated with CheckM (36). Assembly statistics were computed with QUAST (37). * indicate the strains retained by dRep after dereplication.

**Table 2. Functional analysis of the specific genes**. Functional analyses were performed with anvi’o (40), using COGs (41), and Mantis (52). Pathways correspond to COG pathways as indicated by anvi’o. * indicate putative HGT events. Unknown genes correspond to genes without hits.

**Table 3. Differential characteristics between *Snodgrassella alvi* (1. NCIMB 14803**^**T**^**), *Snodgrassella gandavensis* (Bombus group4) (2. LMG 30236**^**T**^, **3. R-53680) and *Snodgrassella communis* (Bombus group5) (4. LMG 28360**^**T**^, **5. R-53528, 6. R-54236)**. +: present, -: absent, w: weak reaction, nd: not determined

**Figure S1. *Neisseriaceae* phylogeny**. The maximum likelihood tree was inferred from 254 core genes under the PROTGAMMALGF model with RAxML (44) from a supermatrix of 111 organisms x 86,654 unambiguously aligned amino-acid positions. Bootstrap support values are shown at the nodes.

**Figure S2. Comparison of the *Snodgrassella* bootstrap phylogeny and the leave-one-out phylogeny**. The maximum likelihood tree was inferred on 254 core genes under the PROTGAMMALGF model with RAxML (44) from a supermatrix of 76 organisms x 86,654 unambiguously aligned amino-acid positions. 100 concatenations of x 70,000-aa long subsamples of the core genes were generated and the corresponding trees computed with RAxML (44). Bootstrap and leave-one-out support values are shown at the nodes. The leave-one-out tree is a cladogram without informative branch lengths.

**Figure S3. Unrooted *Snodgrassella* phylogeny**. The maximum likelihood tree was inferred on 254 core genes under the PROTGAMMALGF model with RAxML (44) from a supermatrix of 75 organisms x 86,654 unambiguously aligned amino-acid positions, after removal of the outgroup. Bootstrap support values are shown at the nodes.

**Figure S4. Gene flow between phylotype genomes of RefSeq bee gut symbionts**. HGT events were inferred with Metachip (55) on the 561 phylotype genomes (75 S*nodgrassella* + 486 other bacterial bee gut phylotypes) present in RefSeq. Bands connect donors and recipients, colors correspond to the donor of the genes.

**Figure S5. Gut bacteria phylogenomic tree, used as species tree for HGT inference**. The maximum likelihood tree was inferred on 76 core genes under the PROTGAMMALGF model with RAxML (44) from a supermatrix of 843 organisms x 12,930 unambiguously aligned amino-acid positions. Bootstrap support values are shown at the nodes.

**Table S1. Information about the 35 *Neisseriaceae* genomes used in the study**.

**Table S2. Information about the 486 bacterial bee gut phylotypes (without *Snodgrassella*) and 247 other bacterial genomes (without the *Neisseriaceae*) used in the study**.

**Table S3. HGT events reported between the 486 phylotype genomes of RefSeq and *Snodgrassella***. Functional analyses were performed with Mantis (52).

## References

1. Engel P, Moran NA. 2013. Functional and evolutionary insights into the simple yet specific gut microbiota of the honeybee from metagenomic analysis. Gut Microbes 4:60–65.

2. Kwong WK, Engel P, Koch H, Moran NA. 2014. Genomics and host specialization of honeybee and bumble bee gut symbionts. PNAS 111:11509–11514.

3. Zheng H, Perreau J, Powell JE, Han B, Zhang Z, Kwong WK, Tringe SG, Moran NA. 2019. Division of labor in honeybee gut microbiota for plant polysaccharide digestion. PNAS 116:25909–25916.

4. Billiet A, Meeus I, Nieuwerburgh FV, Deforce D, Wäckers F, Smagghe G. 2017. Colony contact contributes to the diversity of gut bacteria in bumblebees (Bombus terrestris). Insect Science 24:270–277.

5. Ellegaard KM, Brochet S, Bonilla-Rosso G, Emery O, Glover N, Hadadi N, Jaron KS, Meer JR van der, Robinson-Rechavi M, Sentchilo V, Tagini F, Engel P. 2019. Genomic changes underlying host specialization in the bee gut symbiont Lactobacillus Firm5. Molecular Ecology 28:2224–2237.

6. Ellegaard KM, Engel P. New Reference Genome Sequences for 17 Bacterial Strains of the HoneyBee Gut Microbiota. Microbiology Resource Announcements 7:e00834–18.

7. Guo J, Wu J, Chen Y, Evans JD, Dai R, Luo W, Li J. 2015. Characterization of gut bacteria at different developmental stages of Asian honeybees, Apis cerana. J Invertebr Pathol 127:110–114.

8. Hammer TJ, L. E, Moran NA. 2021. Thermal niches of specialized gut symbionts: the case of social bees. Proceedings of the Royal Society B: Biological Sciences 288:20201480.

9. Kwong WK, Moran NA. 2016. Gut microbial communities of social bees. Nat Rev Microbiol 14:374–384.

10. Kwong WK, Medina LA, Koch H, Sing K-W, Soh EJY, Ascher JS, Jaffé R, Moran NA. 2017. Dynamic microbiome evolution in social bees. Science Advances 3:e1600513.

11. Martinson VG, Moy J, Moran NA. 2012. Establishment of Characteristic Gut Bacteria during Development of the Honeybee Worker. Applied and Environmental Microbiology 78:2830–2840.

12. Meeus I, Parmentier L, Billiet A, Maebe K, Nieuwerburgh FV, Deforce D, Wäckers F, Vandamme P, Smagghe G. 2015. 16S rRNA Amplicon Sequencing Demonstrates that Indoor-Reared Bumblebees (Bombus terrestris) Harbor a Core Subset of Bacteria Normally Associated with the Wild Host. PLOS ONE 10:e0125152.

13. Moran NA, Hansen AK, Powell JE, Sabree ZL. 2012. Distinctive Gut Microbiota of HoneyBees Assessed Using Deep Sampling from Individual Worker Bees. PLOS ONE 7:e36393.

14. Powell JE, Martinson VG, Urban-Mead K, Moran NA. 2014. Routes of Acquisition of the Gut Microbiota of the HoneyBee Apis mellifera. Applied and Environmental Microbiology 80:7378–7387.

15. Praet J, Parmentier A, Schmid-Hempel R, Meeus I, Smagghe G, Vandamme P. 2018. Large-scale cultivation of the bumblebee gut microbiota reveals an underestimated bacterial species diversity capable of pathogen inhibition. Environmental Microbiology 20:214–227.

16. Sabree ZL, Hansen AK, Moran NA. 2012. Independent Studies Using Deep Sequencing Resolve the Same Set of Core Bacterial Species Dominating Gut Communities of HoneyBees. PLOS ONE 7:e41250.

17. Tola YH, Waweru JW, Hurst GDD, Slippers B, Paredes JC. 2020. Characterization of the Kenyan HoneyBee (Apis mellifera) Gut Microbiota: A First Look at Tropical and Sub-Saharan African Bee Associated Microbiomes. 11. Microorganisms 8:1721.

18. Zheng H, Steele MI, Leonard SP, Motta EVS, Moran NA. 2018. Honeybees as models for gut microbiota research. Lab Anim 47:317–325.

19. Martinson VG, Danforth BN, Minckley RL, Rueppell O, Tingek S, Moran NA. 2011. A simple and distinctive microbiota associated with honeybees and bumble bees. Mol Ecol 20:619–628.

20. Zhang Z-J, Huang M-F, Qiu L-F, Song R-H, Zhang Z-X, Ding Y-W, Zhou X, Zhang X, Zheng H. 2021. Diversity and functional analysis of Chinese bumblebee gut microbiota reveal the metabolic niche and antibiotic resistance variation of Gilliamella. Insect Science 28:302–314.

21. Kwong WK, Moran NA. 2013. Cultivation and characterization of the gut symbionts of honeybees and bumble bees: description of Snodgrassella alvi gen. nov., sp. nov., a member of the family Neisseriaceae of the Betaproteobacteria, and Gilliamella apicola gen. nov., sp. nov., a member of Orbaceae fam. nov., Orbales ord. nov., a sister taxon to the order “Enterobacteriales” of the Gammaproteobacteria. Int J Syst Evol Microbiol 63:2008–2018.

22. Engel P, Stepanauskas R, Moran NA. 2014. Hidden Diversity in Honeybee Gut Symbionts Detected by Single-Cell Genomics. PLOS Genetics 10:e1004596.

23. Bosmans L, Pozo MI, Verreth C, Crauwels S, Wilberts L, Sobhy IS, Wäckers F, Jacquemyn H, Lievens B. 2018. Habitat-specific variation in gut microbial communities and pathogen prevalence in bumblebee queens (Bombus terrestris). PLOS ONE 13:e0204612.

24. Horak RD, Leonard SP, Moran NA. 2020. Symbionts shape host innate immunity in honeybees. Proceedings of the Royal Society B: Biological Sciences 287:20201184.

25. Maes PW, Rodrigues PAP, Oliver R, Mott BM, Anderson KE. 2016. Diet-related gut bacterial dysbiosis correlates with impaired development, increased mortality and Nosema disease in the honeybee (Apis mellifera). Molecular Ecology 25:5439–5450.

26. Rothman JA, Leger L, Graystock P, Russell K, McFrederick QS. 2019. The bumble bee microbiome increases survival of bees exposed to selenate toxicity. Environmental Microbiology 21:3417–3429.

27. Sauers LA, Sadd BM. 2019. An interaction between host and microbe genotypes determines colonization success of a key bumble bee gut microbiota member. Evolution 73:2333–2342.

28. Bees of Europe - Hymenoptera of Europe 1 - NAP Editions.

29. Maebe K, Vereecken NJ, Piot N, Reverté S, Cejas D, Michez D, Vandamme P, Smagghe G. 2021. The Holobiont as a Key to the Adaptation and Conservation of Wild Bees in the Anthropocene. Frontiers in Ecology and Evolution 9:849.

30. Moran NA. 2015. Genomics of the honeybee microbiome. Current Opinion in Insect Science 10:22–28.

31. Ludvigsen J, Amdam GV, Rudi K, L’Abée-Lund TM. 2018. Detection and Characterization of Streptomycin Resistance (strA-strB) in a Honeybee Gut Symbiont (Snodgrassella alvi) and the Associated Risk of Antibiotic Resistance Transfer. Microb Ecol 76:588–591.

32. Kwong WK, Moran NA. 2015. Evolution of host specialization in gut microbes: the bee gut as a model. Gut Microbes 6:214–220.

33. Steele MI, Kwong WK, Whiteley M, Moran NA. 2017. Diversification of Type VI Secretion System Toxins Reveals Ancient Antagonism among Bee Gut Microbes. mBio 8:e01630–17.

34. Steele MI, Moran NA. 2021. Evolution of Interbacterial Antagonism in Bee Gut Microbiota Reflects Host and Symbiont Diversification. mSystems 6:e00063–21.

35. Koch H, Abrol DP, Li J, Schmid-Hempel P. 2013. Diversity and evolutionary patterns of bacterial gut associates of corbiculate bees. Molecular Ecology 22:2028–2044.

36. Haft DH, DiCuccio M, Badretdin A, Brover V, Chetvernin V, O’Neill K, Li W, Chitsaz F, Derbyshire MK, Gonzales NR, Gwadz M, Lu F, Marchler GH, Song JS, Thanki N, Yamashita RA, Zheng C, Thibaud-Nissen F, Geer LY, Marchler-Bauer A, Pruitt KD. 2018. RefSeq: an update on prokaryotic genome annotation and curation. Nucleic Acids Research 46:D851–D860.

37. Nasko DJ, Koren S, Phillippy AM, Treangen TJ. 2018. RefSeq database growth influences the accuracy of k-mer-based lowest common ancestor species identification. Genome Biology 19:165.

38. Parks DH, Imelfort M, Skennerton CT, Hugenholtz P, Tyson GW. 2015. CheckM: assessing the quality of microbial genomes recovered from isolates, single cells, and metagenomes. Genome Res 25:1043–1055.

39. Gurevich A, Saveliev V, Vyahhi N, Tesler G. 2013. QUAST: quality assessment tool for genome assemblies. Bioinformatics 29:1072–1075.

40. Bankevich A, Nurk S, Antipov D, Gurevich AA, Dvorkin M, Kulikov AS, Lesin VM, Nikolenko SI, Pham S, Prjibelski AD, Pyshkin AV, Sirotkin AV, Vyahhi N, Tesler G, Alekseyev MA, Pevzner PA. 2012. SPAdes: A New Genome Assembly Algorithm and Its Applications to Single-Cell Sequencing. Journal of Computational Biology 19:455–477.

41. Léonard RR, Leleu M, Vlierberghe MV, Kerff F, Baurain D. 2020. ToRQuEMaDA: Tool for Retrieving Queried Eubacteria, Metadata and Dereplicating Assemblies. bioRxiv 2020.11.15.363259.

42. Eren AM, Esen ÖC, Quince C, Vineis JH, Morrison HG, Sogin ML, Delmont TO. 2015. Anvi’o: an advanced analysis and visualization platform for ‘omics data. PeerJ 3:e1319.

43. Tatusov RL, Galperin MY, Natale DA, Koonin EV. 2000. The COG database: a tool for genome-scale analysis of protein functions and evolution. Nucleic Acids Res 28:33–36.

44. Criscuolo A, Gribaldo S. 2010. BMGE (Block Mapping and Gathering with Entropy): a new software for selection of phylogenetic informative regions from multiple sequence alignments. BMC Evol Biol 10:210.

45. Roure B, Rodriguez-Ezpeleta N, Philippe H. 2007. SCaFoS: a tool for Selection, Concatenation and Fusion of Sequences for phylogenomics. BMC Evolutionary Biology 7:S2.

46. Stamatakis A. 2006. RAxML-VI-HPC: maximum likelihood-based phylogenetic analyses with thousands of taxa and mixed models. Bioinformatics 22:2688–2690.

47. Felsenstein J. 2004. PHYLIP (Phylogeny Inference Package) version 3.6. Distributed by the author. http://www.evolution.gs.washington.edu/phylip.html.

48. Olm MR, Brown CT, Brooks B, Banfield JF. 2017. dRep: a tool for fast and accurate genomic comparisons that enables improved genome recovery from metagenomes through de-replication. 12. The ISME Journal 11:2864–2868.

49. Katoh K, Standley DM. 2013. MAFFT Multiple Sequence Alignment Software Version 7: Improvements in Performance and Usability. Mol Biol Evol 30:772–780.

50. Jain C, Rodriguez-R LM, Phillippy AM, Konstantinidis KT, Aluru S. 2018. High throughput ANI analysis of 90K prokaryotic genomes reveals clear species boundaries. Nat Commun 9:5114.

51. Irisarri I, Baurain D, Brinkmann H, Delsuc F, Sire J-Y, Kupfer A, Petersen J, Jarek M, Meyer A, Vences M, Philippe H. 2017. Phylotranscriptomic consolidation of the jawed vertebrate timetree. Nature Ecology & Evolution 1:1370–1378.

52. Simion P, Philippe H, Baurain D, Jager M, Richter DJ, Di Franco A, Roure B, Satoh N, Quéinnec É, Ereskovsky A, Lapébie P, Corre E, Delsuc F, King N, Wörheide G, Manuel M. 2017. A Large and Consistent Phylogenomic Dataset Supports Sponges as the Sister Group to All Other Animals. Current Biology 27:958–967.

53. Queirós P, Delogu F, Hickl O, May P, Wilmes P. 2021. Mantis: flexible and consensus-driven genome annotation. GigaScience 10.

54. Chaumeil P-A, Mussig AJ, Hugenholtz P, Parks DH. 2020. GTDB-Tk: a toolkit to classify genomes with the Genome Taxonomy Database. Bioinformatics 36:1925–1927.

55. Parks DH, Chuvochina M, Waite DW, Rinke C, Skarshewski A, Chaumeil P-A, Hugenholtz P. 2018. A standardized bacterial taxonomy based on genome phylogeny substantially revises the tree of life. 10. Nature Biotechnology 36:996–1004.

56. Song W, Wemheuer B, Zhang S, Steensen K, Thomas T. 2019. MetaCHIP: community-level horizontal gene transfer identification through the combination of best-match and phylogenetic approaches. Microbiome 7:36.

57. Boratyn GM, Schäffer AA, Agarwala R, Altschul SF, Lipman DJ, Madden TL. 2012. Domain enhanced lookup time accelerated BLAST. Biology Direct 7:12.

58. MacFaddin JF. 2000. Biochemical tests for identification of medical bacteria /. Williams and Wilkins, Baltimore (Md.) :

59. Galtier N. 2007. A Model of Horizontal Gene Transfer and the Bacterial Phylogeny Problem. Systematic Biology 56:633–642.

60. Gribaldo S, Brochier C. 2009. Phylogeny of prokaryotes: does it exist and why should we care? Research in Microbiology 160:513–521.

61. Stanhope MJ, Lupas A, Italia MJ, Koretke KK, Volker C, Brown JR. 2001. Phylogenetic analyses do not support horizontal gene transfers from bacteria to vertebrates. Nature 411:940–944.

62. Siu-Ting K, Torres-Sánchez M, San Mauro D, Wilcockson D, Wilkinson M, Pisani D, O’Connell MJ, Creevey CJ. 2019. Inadvertent Paralog Inclusion Drives Artifactual Topologies and Timetree Estimates in Phylogenomics. Molecular Biology and Evolution 36:1344–1356.

63. Struck TH. 2013. The Impact of Paralogy on Phylogenomic Studies – A Case Study on Annelid Relationships. PLOS ONE 8:e62892.

64. Li Y, Xue H, Sang S, Lin C, Wang X. 2017. Phylogenetic analysis of family Neisseriaceae based on genome sequences and description of Populibacter corticis gen. nov., sp. nov., a member of the family Neisseriaceae, isolated from symptomatic bark of Populus × euramericana canker. PLOS ONE 12:e0174506.

65. Powell E, Ratnayeke N, Moran NA. 2016. Strain diversity and host specificity in a specialized gut symbiont of honeybees and bumblebees. Molecular Ecology 25:4461–4471.

66. Tindall BJ, Rosselló-Móra R, Busse H-J, Ludwig W, Kämpfer PY 2010. Notes on the characterization of prokaryote strains for taxonomic purposes. International Journal of Systematic and Evolutionary Microbiology 60:249–266.

67. Goris J, Konstantinidis KT, Klappenbach JA, Coenye T, Vandamme P, Tiedje JMY 2007. DNA–DNA hybridization values and their relationship to whole-genome sequence similarities. International Journal of Systematic and Evolutionary Microbiology 57:81–91.

68. Richter M, Rosselló-Móra R. 2009. Shifting the genomic gold standard for the prokaryotic species definition. PNAS 106:19126–19131.

69. Dong Z-X, Li H-Y, Chen Y-F, Wang F, Deng X-Y, Lin L-B, Zhang Q-L, Li J-L, Guo J. 2020. Colonization of the gut microbiota of honeybee (Apis mellifera) workers at different developmental stages. Microbiological Research 231:126370.

70. Dehon M, Engel MS, Gérard M, Aytekin AM, Ghisbain G, Williams PH, Rasmont P, Michez D. 2019. Morphometric analysis of fossil bumble bees (Hymenoptera, Apidae, Bombini) reveals their taxonomic affinities. ZooKeys 891:71–118.

71. Hines HM. 2008. Historical biogeography, divergence times, and diversification patterns of bumble bees (Hymenoptera: Apidae: Bombus). Syst Biol 57:58–75.

72. Williams PH, Huang J, Rasmont P, An J. 2016. <p><strong>Early-diverging bumblebees from across the roof of the world: the high-mountain subgenus <em>Mendacibombus</em> revised from species’ gene coalescents and morphology (Hymenoptera, Apidae)</strong></p>. 1. Zootaxa 4204:1–72.

73. Cardinal S, Straka J, Danforth BN. 2010. Comprehensive phylogeny of apid bees reveals the evolutionary origins and antiquity of cleptoparasitism. PNAS 107:16207–16211.

74. Storz G, Wolf YI, Ramamurthi KS. 2014. Small Proteins Can No Longer Be Ignored. Annual Review of Biochemistry 83:753–777.

75. Lluch-Sena M, Delgado J, Chen W, Llorens V, O’Reilly F, Wodke JA, Unal B, Yus E, Martinez S, Nichols RJ, Vivancos A, Schmeisky A, Stulke J, Van Noort V, Gavin A, Bork P, Serrano L. 2015. Defining a minimal cell: essentiality of small ORFs and ncRNAs in a genome-reduced bacterium. Molecular Systems Biology 11:780.

76. Duval M, Cossart P. 2017. Small bacterial and phagic proteins: an updated view on a rapidly moving field. Current Opinion in Microbiology 39:81–88.

77. Thomas AM, Segata N. 2019. Multiple levels of the unknown in microbiome research. BMC Biology 17:48.

78. The UniProt Consortium. 2021. UniProt: the universal protein knowledgebase in 2021. Nucleic Acids Research 49:D480–D489.

79. Powell JE, Leonard SP, Kwong WK, Engel P, Moran NA. 2016. Genome-wide screen identifies host colonization determinants in a bacterial gut symbiont. PNAS 113:13887–13892.

