## Supplementary figures and images for "Phylogenomic analyses of *Snodgrassella* isolates from honeybees and bumblebees reveals taxonomic and functional diversity"

### FigureS1

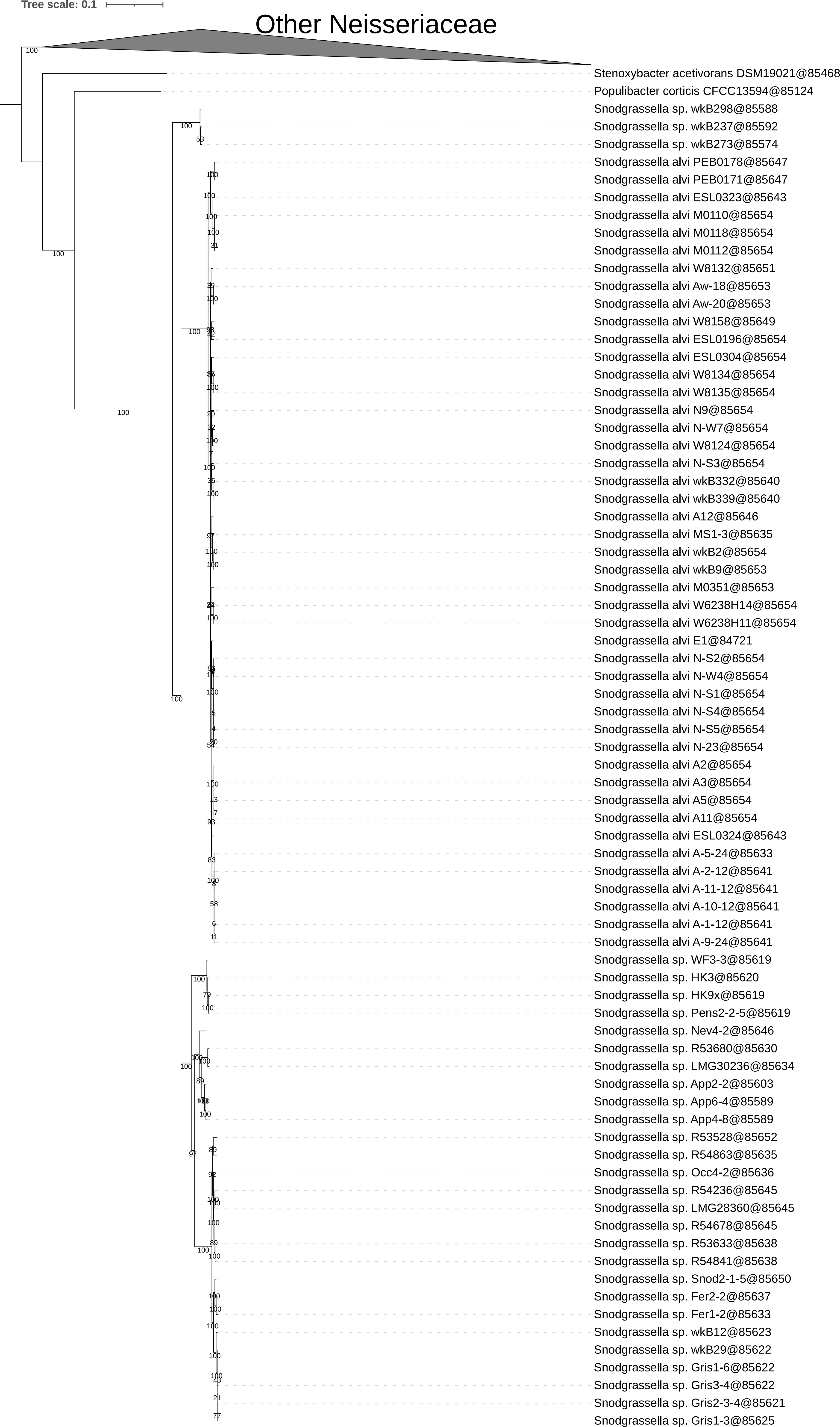

### FigureS2

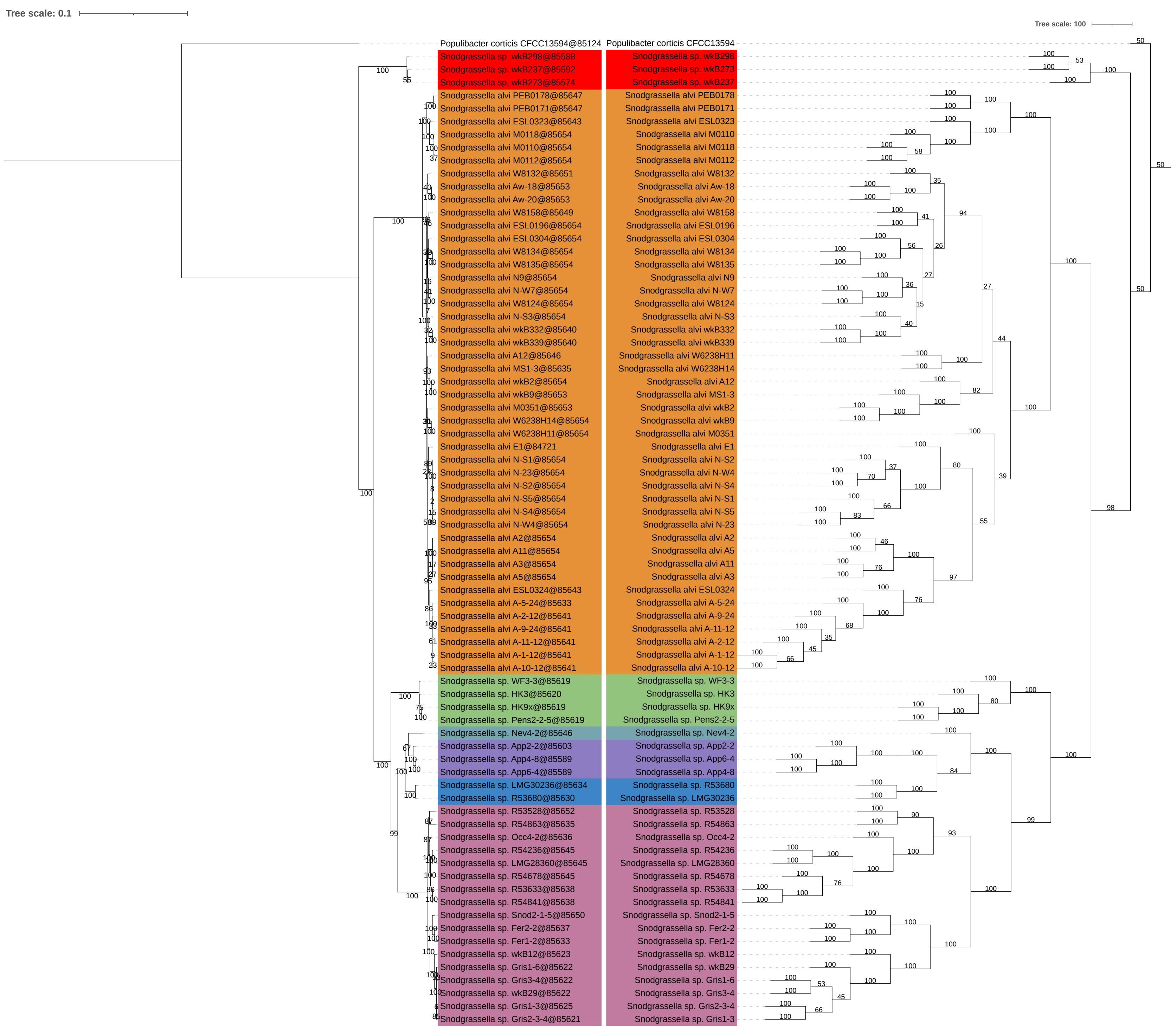

### FigureS3

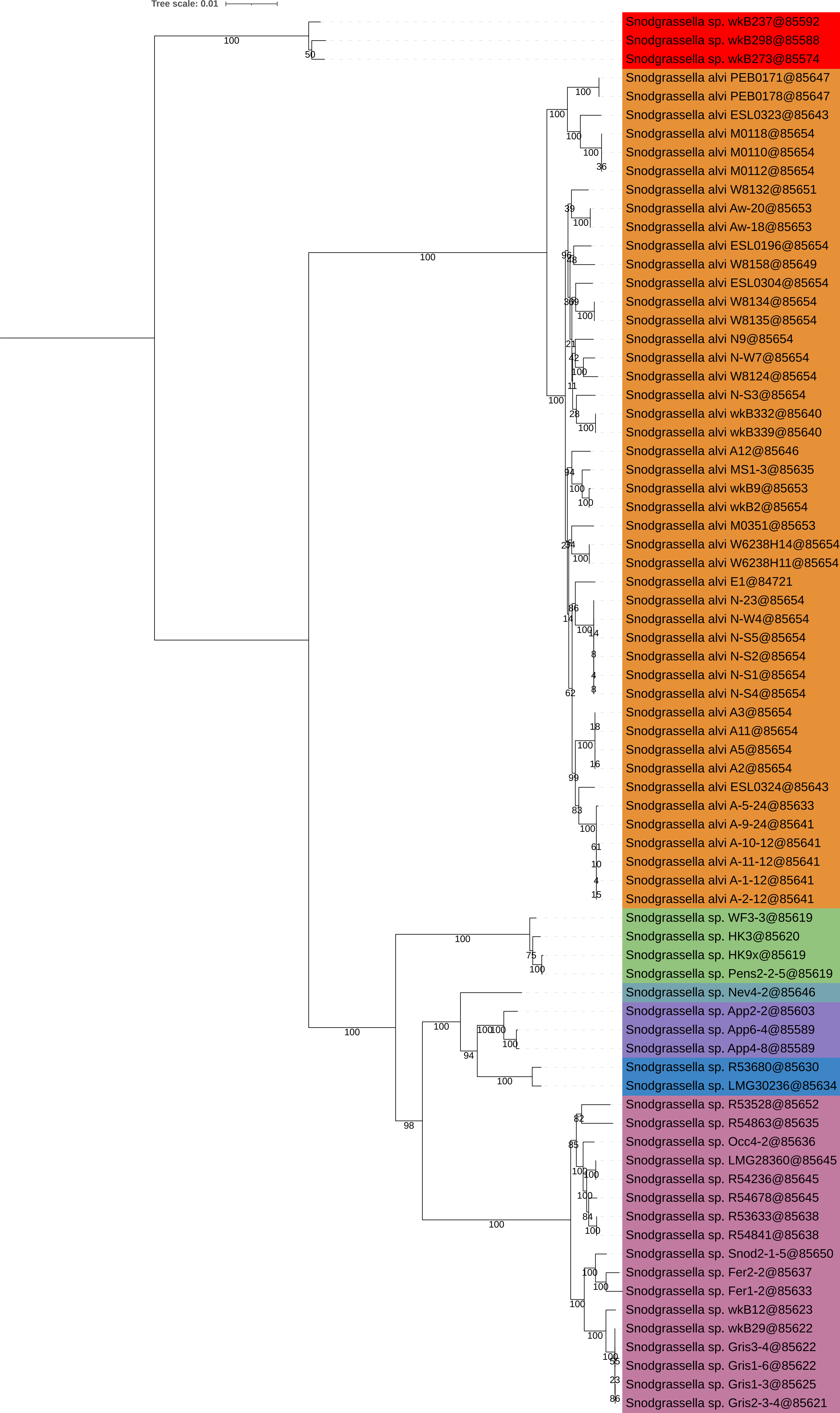

### FigureS4

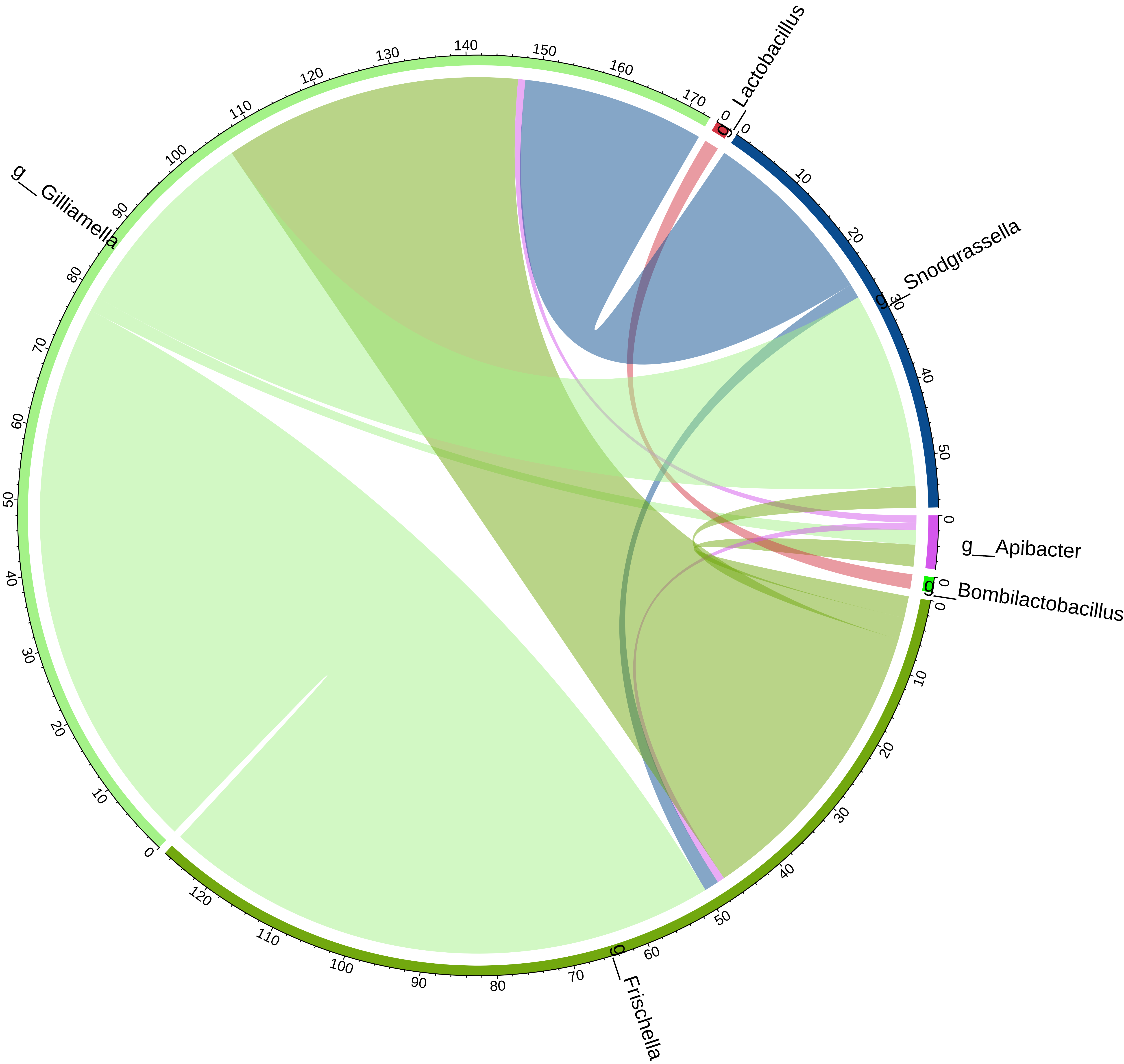

### FigureS5

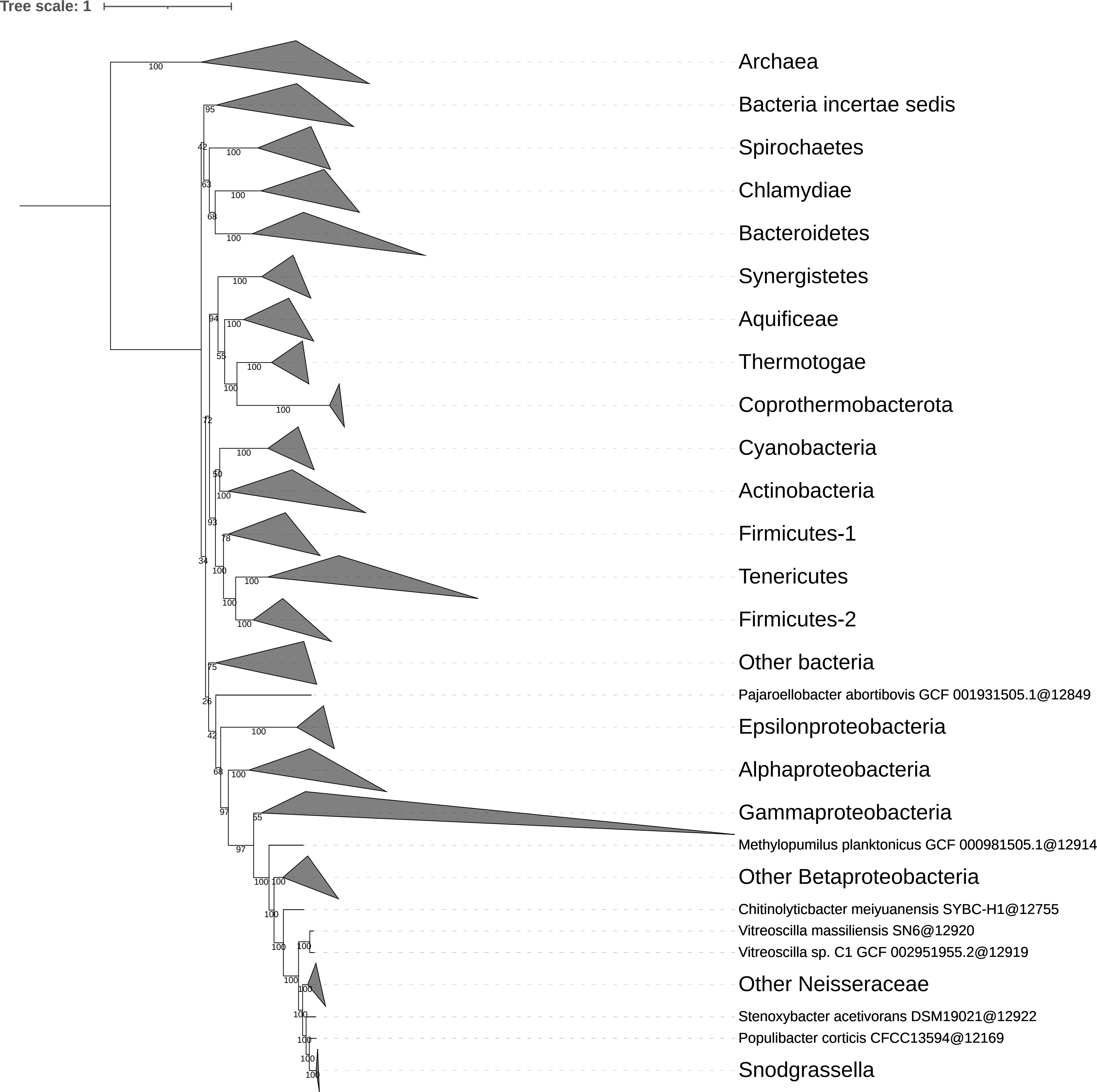
