## Supplementary material for "Phylogenomic analyses of *Snodgrassella* isolates from honeybees and bumblebees reveals taxonomic and functional diversity": Supp-note

**Supplementary information: detailed species descriptions.**

### Description of *Snodgrassella gandavensis* sp. nov

*Snodgrassella gandavensis* (gan.da.ven'sis. M.L. masc. adj. *gandavensis* of *Gandavum*, the Latin name for Ghent, referring to the place where these bacteria were first isolated).

Cells are non-motile, Gram stain-negative rods, about 1.2 µm long and 0.8 µm wide that occur singly or in pairs. Optimal growth on AC agar at 37°C in a $CO_2$ enriched atmosphere. After two days of incubation using these conditions, colonies are beige, round, smooth, shiny, translucent and about 0.5 mm in diameter. Growth also occurs at 40°C and weakly at 28°C, but not at 4, 15 and 20 °C. At 37°C in a $CO_2$ enriched atmosphere, growth also occurs on nutrient agar (LMG 30236$^T$ weakly), but not on BHI agar. Growth on TSA and Columbia agar with 5% sheep blood is strain dependent. Grows at 37°C on AC agar and AC agar supplemented with 10 mM $KNO_3$ in anaerobic conditions, but not in aerobic conditions.

Grows on AC agar in the presence of 0-10% (w/v) NaCl, although weakly at ≥ 2.0% NaCl. Grows optimally on AC agar at pH 6 and weakly at pH 7. Strain dependent growth at pH 4, 5, 8 and 9. Positive for catalase activity, nitrate reduction, urease, indole production, growth on AC agar with 0.8% gelatin, AC agar with starch, AC agar with skim milk and AC agar with tween 20 and tween 80, hydrolysis of tween 20 and 80. Negative for oxidase activity, denitrification, H2S production, gelatinase activity, growth on DNase agar, hydrolysis of starch and casein.
The type strain is LMG 30236$^T$ (= CECT 30450$^T$), which was isolated in 2015 from the gut of *Bombus pascuorum* sampled in 2015 in Wetteren, Belgium. The whole genome sequence of LMG 30236$^T$ has a size of 2.51 Mbp. The DNA G+C content is 43.76 mol%. The whole genome sequence is publicly available under accession number GCA_914768025.1. The 16S rRNA gene sequence is publicly available under accession number OU943324.

### Description of *Snodgrassella communis* sp. nov.

*Snodgrassella communis* (com.mu'nis. L. fem. adj. *communis* common, because of its wide host range).

Cells are non-motile, Gram stain-negative rods, about 1.2 µm long and 0.8 µm wide that occur singly or in pairs.
Growth on AC agar, BHI agar and Columbia agar with 5% sheep blood at 37°C in a $CO_2$ enriched atmosphere and on nutrient agar and TSA, although the latter growth can be weak depending on the strain. After two days of incubation on AC agar in a $CO_2$ enriched atmosphere, colonies are beige, round with an irregular margin, translucent and about 0.5 mm in diameter. Growth also occurs at 40°C and weakly at 28°C, but not at 4, 15, 20 and 45 °C. Grows at 37°C on AC agar and AC agar supplemented with 10 mM $KNO_3$ in anaerobic conditions, growth in aerobic conditions is strain dependent. Grows on AC agar in the presence of 0-10% (w/v) NaCl, although weakly at ≥ 2.0-3.0 % NaCl. Grows optimally on AC agar at pH 6, and weak at pH 4 and 5. Strain dependent growth at pH 7, 8 and 9. Positive for catalase activity, nitrate reduction, indole production, growth on AC agar with 0.8% gelatin, AC agar with starch, AC agar with skim milk and AC agar with tween 20 and tween 80. Negative for oxidase activity, haemolysis, denitrification, urease, H2S production, gelatinase activity, growth on DNase agar, hydrolysis of starch, casein and tween 20. Strain dependent hydrolysis of tween 80.

The type strain is LMG 28360$^T$ (= CECT 30451$^T$), which was isolated in 2013 from the gut of *Bombus terrestris* sampled in 2013 in Ghent, Belgium. The whole genome sequence of LMG 28360$^T$ has a size of 2.31 Mbp. The DNA G+C content is 43.26 mol%. The whole genome sequence is publicly available under accession number GCA_914068745.1. The 16S rRNA gene sequence is publicly available under accession number OU943323.
